# Multimodal profiling of term human decidua reveals tissue-specific immune adaptations with maternal obesity

**DOI:** 10.1101/2022.11.01.514598

**Authors:** Suhas Sureshchandra, Brianna M. Doratt, Norma Mendoza, Monica Rincon, Nicole E. Marshall, Ilhem Messaoudi

**Author notes:** Corresponding Author: Ilhem Messaoudi, Department of Microbiology, Immunology and Molecular Genetics, College of Medicine, University of Kentucky, 760 Press Avenue, Lexington, KY 40536.

## Abstract

The leukocyte diversity of early maternal-fetal interface has been recently described, however, characterization of decidua at term is still pending. We, therefore, profiled the CD45+ compartment within human term decidua collected via cesarean section. Relative to the first trimester, our analyses revealed a shift from NK cells and macrophages to T-cells and enhanced activation of all immune cells. T-cells in the decidua were phenotypically distinct from their blood counterparts and demonstrated significant clonotype sharing. Using pseudo-temporal analyses of macrophages, we characterize previously undescribed diversity of bonafide tissue resident decidual macrophages. The frequency of this population correlated with high pregravid maternal body mass index but responded poorly to bacteria ligands. Collectively, these findings provide evidence of skewing of decidual myeloid cells towards immunoregulation representing a possible mechanism to safeguard the fetus. This immune cell atlas of term decidua is an asset for future studies investigating pathological conditions that compromise reproductive success.

**One sentence summary:** Human immune cell atlas of term decidua and observed alterations with pre-gravid obesity.

## INTRODUCTION

The placenta is a complex organ composed of maternal derived membranes -the decidua- and membranes of fetal origin -the chorionic villi ^1^. Functionally, these membranes facilitate gas, nutrient, and waste exchanges between the fetus and the mother. Moreover, the placental membranes harbor a unique immune landscape, which facilitates trophoblast invasion, maternal tolerance, debris clearance, and anti-microbial defense ^2^. Longitudinal changes in frequencies, phenotype, and functions of decidual immune cells with healthy gestation are of great interest, given the role of the placenta in the pathophysiology of several obstetric complications such as preterm labor ^3^, preeclampsia ^4–6^ and chorioamnionitis ^7^. First-trimester decidua (10-12 weeks) is comprised mostly of decidual Natural Killer cells (dNKs, ~70% of immune cells), decidual macrophages (20-25%), and T-cells (3-10%) ^8, 9^. The dNKs are the earliest invading immune cells in the decidua, playing a critical role in regulating trophoblast invasion and decidualization ^10^. Decidual macrophages and T-cells play important roles in maintaining immune tolerance ^11^, anti-microbial defense ^2^, spiral artery remodeling ^12^, and clearance of apoptotic bodies from the placental vasculature ^13^. While the diversity and characteristics of decidual immune cells during the first trimester have been well described, those at term remain poorly defined. Over the course of healthy gestation, current models predict that dNK and macrophage numbers decrease while those of T-cells remain intact in the decidua basalis ^14^. However, there is still no consensus about the phenotypic and functional diversity of decidual T-cells and macrophages at term.

Maternal pre-pregnancy (pregravid) obesity is associated with an increased risk for several obstetric complications including gestational diabetes and gestational hypertension ^15, 16^, preeclampsia ^17^, placental abruption ^18^, preterm delivery ^19, 20^, and need for cesarean section ^21, 22^. These predispositions are mediated by altered placental structure ^23, 24^, increased levels of oxidative stress ^25^, and elevated expression of pro-inflammatory factors ^26^. Studies in animal models suggest that changes in the decidual inflammatory environment can be attributed to dysregulated cell turnover and immune cell rewiring ^27, 28^, however, these changes have been poorly defined in humans. For example, first-trimester uterine NK cells in mothers with obesity exhibit an imbalance of inhibitory and activating receptors resulting in aberrant activation ^29^. Immunohistochemistry and flow cytometry-based studies have reported increased CD68+ macrophage frequencies in chorionic villi with maternal obesity ^30^. In line with these observations, higher transcript levels of pro-inflammatory genes IL6, CCL2, IL8, and TLR4 were detected in a baboon model of HFD-induced obesity ^31, 32^, suggesting potential immune cell adaptations with obesity at the maternal-fetal interface. However, a clear consensus on these observations is lacking, perhaps due to differences in choices of placental membranes analyzed, immune markers used, experimental readouts measured, and confounding variables in human studies ^23^.

A detailed understanding of immune cell adaptations in the decidua that support late pregnancy is currently lacking. More importantly, how maternal obesity reprograms the immune milieu of decidua at term is yet to be fully assessed. To address these gaps in our knowledge, we carried out a multi-pronged study using single-cell RNA sequencing, flow cytometry, luminex, and functional assays to profile the diverse subsets of macrophages, T-cells, and NK cells in term decidua and compared them to first-trimester decidua atlas ^9^. More importantly, using placental decidua isolated from a cohort of pregnant mothers undergoing cesarean in the absence of any obstetric complications, we studied the impact of pregravid obesity on immune cell adaptations at term.

Our data show that, relative to first trimester, decidual immune cells exhibit signs of heightened activation status in healthy pregnancies. Furthermore, the relative abundance of T-cells increases at term with a concomitant reduction in the frequency of macrophages and NK cells. We also report a shift from the dNK1 subset at 12 weeks to dNK3 subset at term. T-cell production of pro-inflammatory cytokines was reduced while production of regulatory and Th17 cytokines increased in decidual T-cells when compared to circulating T-cells, in line with immune tolerance. Finally, we observed two major macrophage subsets that can be distinguished based on HLA-DR and CD11c expression and may represent a tissue-resident and a monocyte-derived subset. Pregravid obesity led to a reduction of T-cell numbers but increased accumulation of macrophages and functional rewiring of HLA-DR^high^CD11c^high^ macrophages. Furthermore, decidual macrophages from women with obesity generated dampened responses to bacterial ligands, suggesting skewing towards regulatory phenotype. Collectively, our study provides an atlas of the immune landscape of term decidua and shifts mediated by pregravid obesity.

## METHODS

### Human subjects

This study was approved by the Institutional Ethics Review Board of Oregon Health & Science University and the University of California, Irvine. A total of 102 non-smoking women (47 lean, 12 overweight, and 43 obese) who had an uncomplicated pregnancy were enrolled for this study. Written informed consent was obtained from all participants. Due to the strong positive correlation between pre-pregnancy BMI and total body fat in our previous studies ^33^, we used pre-pregnancy BMI as a surrogate for maternal pregravid obesity and stratified participants into lean or obese based on pre-pregnancy BMI (Table 1). Only samples from term cesarean deliveries were included in the analysis. Exclusion criteria include active maternal infection, documented fetal congenital anomalies, substance abuse, chronic illness requiring regular medication use, preeclampsia, gestational diabetes, chorioamnionitis, and significant medical conditions (active cancers, cardiac, renal, hepatic, or pulmonary diseases), or an abnormal glucose tolerance test.

**Table 1:** Cohort Characteristics. Avg (SD). *p<=0.0001

|  | Lean | Overweight | Obese |
| --- | --- | --- | --- |
| Enrollment | 47 | 12 | 43 |
| BMI (kg/m<sup>2</sup>) * | 21.9 (1.7) | 27.4 (1.6) | 38 (7.9) |
| Maternal Age (years) | 33.7 (4.6) | 34.7 (4.71) | 32.4 (4.5) |
| Gestational Age<br>(weeks) | 38.8 (0.7) | 39.02 (0.7) | 38.3 (0.9) |
| % Female | 55.8 | 33.3 | 33.3 |

### Sample collection and processing

Decidua basalis membranes were separated immediately and placed in individual 50 mL conicals immersed in RPMI supplemented with 10% FBS and antibiotics. Samples were collected by the Maternal Fetal Medicine unit at Oregon Health & Science University, Portland OR, and then shipped overnight for further processing. Tissues were washed in HBSS to remove contaminating blood and any visible blood vessels macroscopically separated and then washed for 10 minutes in HBSS before processing. Tissues were minced into approximately 0.2-0.3 mm^3^ cubes and enzymatically digested in 0.5mg/mL collagenase V (Sigma, C-9722) solution in 50 mL R3 media (RPMI 1640 with 3% FBS, 1% Penicillin-Streptomycin, 1% L-glutamine, and 1M HEPES) at 37°C for 1 hour. The disaggregated cell suspension was passed through cell strainers to eliminate visible chunks of fat/tissue. Cells were pelleted from the filtrate, passed through a 70-um cell sieve, centrifuged, and resuspended in R3 media. Red blood cells were lysed using RBS lysis buffer (155 mM NH_4_Cl, 12 mM NaHCO_3_, 0.1 mM EDTA in double-distilled water) and resuspended in 5 mL R3. The cell suspension was then layered on a discontinuous 60% and 40% percoll gradient and centrifuged for 60 minutes with brakes off. Immune cells at the interface of 40% and 60% gradients were collected, washed in HBSS, counted, and cryopreserved for future analysis.

### Blood processing

Peripheral blood collected before cesarean was layered on a Ficoll-Paque (GE Healthcare) gradient and centrifuged at 2000 rpm for 30 minutes without brakes. Peripheral Blood Mononuclear cells (PBMC) were resuspended in FetalPlex (Gemini BioProducts) with 10% DMSO (Sigma) and stored in liquid nitrogen for future analysis. Plasma samples were stored at −80°C until they were used.

### Immunophenotyping

For broad phenotyping of immune cells within term decidua, 1-2 X 10^6^ fresh decidual leukocytes were washed with PBS and stained using the following cocktail of antibodies: CD45, CD20, CD4, CD8b, CD14, HLA-DR, CD56, CD16, CD11c, CD123 (Biolegend, San Diego, CA), for 30 minutes in the dark at 4°C.

For T-cell phenotyping, 0.5 X 10^6^ freshly thawed decidual leukocytes were stained using the following panel of surface antibodies: CD4, CD8, CD45RA, CCR7, CD69, CD103, CTLA4, CD25, and PD-1 (Biolegend, San Diego, CA) for 30 minutes at 4°C. Cells were then fixed and permeabilized for 30 minutes followed by intra-nuclear staining for FOXP3 for an additional 4hours using FOXP3 Fix/Perm Buffer set (Biolegend, San Diego, CA).

For identification of NK cell subsets, 0.5 X 10^6^ freshly thawed decidual leukocytes were stained using the following panel of antibodies: CD56, CD16, CD103, CD49a, CD39, ITGB2, KIR2DL4, NKG2A, NKG2C, and CD9 (Biolegend, San Diego, CA).

For a more comprehensive analysis of macrophage phenotypes, 0.5 X 10^6^ thawed decidual cells were stained using the following panel of antibodies CD14, HLA-DR, CD11c, CD16, CD68, CD86, TLR4, CD163, CD206, and CD209. A second aliquot of cells was surface stained with CD14, HLA-DR, TREM1, TREM2, CD64, GLUT1, CD9, FOLR2, CCR2 (Biolegend, San Diego, CA), followed by fixation/permeabilization before staining intracellularly with TREM2 and S100A9 for 4 hours.

All surface staining was done in the presence of Human Fc block and Monocyte Blocker (Biolegend, San Diego, CA). Following staining, all samples were washed twice in FACS buffer and acquired with the Attune NxT Flow Cytometer (ThermoFisher Scientific, Waltham, MA), immediately after the addition of SYTOX Live Dead Cell Stain (ThermoFisher Scientific) for unfixed samples. Data were analyzed using FlowJo 10.5 (Beckton Dickinson, Ashland, OR).

### Ex vivo stimulation and intracellular cytokine staining

To measure cytokine responses of decidua macrophages, 10^6^ thawed cells were stimulated for 6h at 37°C in RPMI supplemented with 10% FBS in the presence or absence of bacterial TLR cocktail containing 1 ug/mL LPS (TLR4 ligand, E.coli 055:B5; Invivogen, San Diego, CA), 2 ug/mL Pam3CSK4 (TLR1/2 agonist, Invivogen), and 1 ug/mL FSL-1 (TLR2/6 agonist, Sigma-Aldrich), or 6 X 10^5^ cfu/mL E. coli (Escherichia coli (Migula) Castellani and Chalmers ATCC 11775).

For NK and T-cell responses, individual aliquots of 10^6^ thawed cells were stimulated for 4h at 37°C in RPMI supplemented with 10% FBS in the presence or absence of 500 ng PMA and 1 ug Ionomycin (Sigma Aldrich). For efficient capture of NK cell degranulation, CD107a was added during stimulation.

For all samples, Brefeldin A (Sigma, St. Louis, MO) was added after 1 -hour incubation and cell were cultured for an additional 5 hours before surface and intracellular cytokine staining.

Cells were stained with the following surface antibodies: [1] CD14 and HLA-DR for macrophages; [2] CD4 and CD8 for T-cells; and [3] CD56, CD16, CD103, CD49a, CD39, and ITGB2 for NK cells for 30 min in the dark at 4°C. Samples were fixed and permeabilized using Fixation buffer (Biolegend) at 4°C for 20 min and stained intracellularly overnight for TNFα and IL-6, and IL-1b for macrophages, IFNg, TNFa, IL-2, IL-4, IL-17, TGF-b and MIP-1b for T-cells and IFNg, GZMB, and MIP-1b for NK cells. Cells were then washed and acquired using the Attune NxT Flow Cytometer (ThermoFisher Scientific, Waltham MA) and analyzed using FlowJo 10.5 (BD, Ashland OR).

### Imaging flow cytometry

For visualization of macrophage subsets and nuclear translocation of NK-kB p50, 500,000 decidual leukocytes were washed and stained for dead cell exclusion (Ghost Violet 510, 1:1000 dilution) for 30 minutes at 4°C. Cells were then surface stained (CD14, CD11c, HLA-DR) and fixed with 1X fixation buffer (Luminex Corporation, Austin, TX) for 10 minutes at room temperature (RT) before the addition of p50-AF488 (1:20 dilution) in 1X Assay Buffer for 30 minutes at RT in the dark. At the end of the incubation, cells were washed twice in 1X Assay buffer and resuspended in 50 uL 0.25X Fixation Buffer in polypropylene Eppendorf tubes. After the addition of nuclear dye (7-AAD, 1:50), samples were run on Amnis ImageStream XMark II Imaging flow cytometer (Luminex Corporation) and data were analyzed on Ideas Analysis Software (Luminex Corporation).

### Phospho Flow

Decidual leukocytes were washed with FACS buffer and surface stained for CD14 and HLA-DR in FACS tubes. Pellets were washed in FACS buffer and resuspended in 100 uL prewarmed PBS (Ca+ Mg+ free). Cells were fixed immediately by the addition of equal volumes of prewarmed Cytofix Buffer (BD Biosciences) and thorough mixing and incubating at 37C for 10 minutes. Cells were then centrifuged at 600g for 8 minutes. Supernatants were removed leaving no more than 50 uL residual volume. Cells were then permeabilized by the addition of 1 mL 1X BD PermWash Buffer I, mixed well, and incubated at RT for 30 minutes. Pellets were then spun, aspirated, and stained intranuclearly with antibodies against NF-kB p65 (pS529) AF647 (Clone K10-895.12.50, Cell Signaling Technology) or IkBa PE (Clone MFRDTRK, eBioscience, San Diego, CA) and Phospho-p38 MAPK-APC (Clone 4NIT4KK, eBioscience) for 1h at RT in the dark. Samples were washed twice in Permwash Buffer I, resuspended in FACS buffer and acquired on the Attune NxT flow cytometer.

### BODIPY Staining

Briefly, 500,000 decidual leukocytes were surface stained (CD14, HLA-DR) for 20 minutes at 4°C, washed twice, and resuspended in 500 uL warm 1X PBS containing 1 ug/mL BODIPY™ 493/503 (ThermoFisher Scientific). Cells were incubated at 37°C for 10 minutes and acquired on the Attune NxT flow cytometer.

### Phagocytosis Assay

500,000 decidual leukocytes were incubated for 2 hours at 37°C in media containing pH-sensitive pHrodo® E. coli BioParticles® conjugates (ThermoFisher Scientific) (1 mg/mL concentration). Pellets were washed twice, surface stained (CD14, HLA-DR) and resuspended in ice-cold FACS buffer. Samples were analyzed using flow cytometry with no pHrodo controls.

### Cytosolic ROS assay

500,000 decidual leukocytes were incubated with 2.5 uM CellROX Deep Red (Life Technology, Carlsbad, CA) at 37°C for 30 minutes. For negative control, cells were incubated in serum-free media containing 200 uM antioxidant N-acetylcysteine (NAC) for 1.5 hours. Both negative and positive controls were incubated with tert-butyl hydroperoxide (TBHP) for 30 minutes to induce oxidative stress. All samples were then surface stained (CD14, HLA-DR) and analyzed using flow cytometry.

### Glucose Uptake Assay

500,000 decidual leukocytes were resuspended in glucose-free media and incubated with 60 mM NBDG (ThermoFisher), a fluorescent glucose analog for 30 minutes at 37°C. Samples were washed in 1X PBS, surface stained (CD14, HLA-DR), and median fluorescence of NBDG recorded using flow cytometry.

### MitoTracker Analysis

500,000 decidual leukocytes were resuspended in 100 uL of 1X MitoTracker Red CMXRos (ThermoFisher) (1:1000 of stock) in 96 well plates, incubated at 37°C for 30 minutes, washed, surface stained (CD14, HLA-DR), and median fluorescence of MitoTracker recorded using flow cytometry.

### Seahorse Assay

Oxygen Consumption Rate (OCR) and Extracellular Acidification Rate (ECAR) were measured using Seahorse XF Cell Energy Phenotype kit on Seahorse XFp Flux Analyzer (Agilent Technologies, Santa Clara, CA) following the manufacturer’s instructions. Briefly, 200,000 sorted macrophage populations (pooled n=3/group) were seeded on Cell-Tak (Corning, Corning, NY) coated 8-well culture plates in phenol-free RPMI media containing 2 mM L-glutamine, 10 mM L-glucose, and 1mM sodium pyruvate. Seeded plates were placed in 37°C incubator without CO_2_ then run on the XFp with extended basal measurements, followed by injection of a stressor cocktail of 1 ug/mL LPS for 1 hour in a 37°C incubator without CO_2_. Plates were run on the XFp for 8 cycles of basal measurements, followed by acute injection of L-glucose (100 mM), oligomycin (ATP synthase inhibitor) carbonilcyanide p-triflouromethoxyphenylhydrazone (FCCP; a mitochondrial uncoupling agent).

### Cytokine production

Live decidua macrophage subsets and peripheral blood monocytes (CD14+ HLA-DR^high^ and HLA-DR^low^, and SYTOX Blue negative) were sorted using BD FACS Aria Fusion. 50,000-sorted cells were placed in a 96-well plate overnight to measure cytokine production. Supernatants were collected and stored at −80°C until further analysis using Luminex. Briefly, immune mediators in cell supernatants were measured using Human Custom 29-plex Multi-Analyte Kit (R&D Systems, Minneapolis, Minnesota) measuring the following analytes: CCL2 (MCP-1), CCL4 (MIP-1b), CXCL9 (MIG), CXCL11 (I-TAC), GM-CSF, IFNg, IL-1RA, IL-4, IL-7, IL-12p70, IL-18, PD-L1, S100B, VEGF, IL-15, CCL3, CCL11, CXCL10, CXCL13, IFNb, IL-1b, IL-2, IL-6, CXCL8 (IL-8), IL-17A, IL-23, PDGF-BB, TNFa, and IL-10. Samples were processed per manufacturer’s instructions, recorded, and analyzed on the Magpix xPONENT system version 4.2 (Luminex Corporation)

### FACS and 3’ Single Cell RNA library preparation

Freshly thawed immune cells from decidua and peripheral blood were stained with CD45-FITC at 4°C in 1% FBS in DPBS without calcium and magnesium. Cells were washed twice and sorted on BD FACS Aria Fusion into RPMI (supplemented with 30% FBS) following the addition of SYTOX Blue stain for dead cell exclusion. Cells were then counted in triplicates on a TC20 Automated Cell Counter (BioRad, Hercules, CA), washed, and resuspended in PBS with 0.04% BSA in a final concentration of 1200 cells/uL. Single-cell suspensions were then immediately loaded on the 10x Genomics Chromium Controller with a loading target of 17,600 cells. Libraries were generated using the V3 chemistry per the manufacturer’s instructions (10x Genomics, Pleasanton CA). Libraries were sequenced on Illumina HiSeq with a sequencing target of 50,000 reads per cell.

### 3’ Single cell RNA-Seq data analysis

Raw reads were aligned and quantified using the Cell Ranger Single-Cell Software Suite (version 3.0.1, 10x Genomics) against the GRCh38 human reference genome internally running the STAR aligner. Downstream processing of aligned reads was performed using Seurat (version 3.1.1). Droplets with ambient RNA (cells with fewer than 400 detected genes), potential doublets (cells with more than 4000 detected genes and dying cells (cells with more than 20% total mitochondrial gene expression) were excluded during initial QC. Data objects from the lean and obese groups were integrated using Seurat ^34^. Data normalization and variance stabilization were performed using SCTransform function ^35^ using a regularized negative binomial regression, correcting for differential effects of mitochondrial and ribosomal gene expression levels and cell cycle.

We integrated placental leukocytes scRNA-Seq datasets with those from matched maternal PBMC using Seurat’s IntegrateData function to remove potential contaminating blood cells. Placenta cells clustering with PBMC were removed from downstream analyses.

Dimensional reduction was performed using RunPCA function to obtain the first 30 principal components followed by clustering using the FindClusters function in Seurat. Clusters were visualized using the UMAP algorithm as implemented by Seurat’s runUMAP function. Cell types were assigned to individual clusters using FindMarkers function with a fold change cutoff of at least 0.4 (FDR ≤ 0.05) and using a known catalog of well-characterized scRNA markers for human PBMC ^36^, tissue-resident lymphoid cells ^9^, and leukocytes in first-trimester human placentas ^9^.

To study the evolution of the immune landscape of placental decidua with gestational age, we used the recently described atlas of first trimester decidual CD45+ cells ^9^. Raw data were downloaded for 6 individuals (D6-10, D12) and analyzed as described above resulting in 24,849 cells. Cells were downsampled (6,116) to match the number of cells from term decidua and integrated using Seurat’s IntegrateData function and markers compared using FindMarkers function.

Temporal trajectory was predicted using Monocle (version 2.8.0). Briefly, t-SNE was used for cell clustering. Monocle’s differentialGeneTest function was used to identify differential genes. Differential genes with a q-value less then 1e-10 were used to ploT-cells in a pseudotime scale. Metascape was used to identify Gene ontology (GO) terms ^37^.

### 5’ GEX and scTCR library preparation

Matched PBMC and decidual leukocytes were thawed, washed, filtered, and stained with Ghost Violet 540 live-dead stain (Tonbo Biosciences, San Diego, CA) for 30 min in the dark at 4°C. Given the parallel assessment of both TCR and gene expression, we used TotalSeq-C, an antibody-based multiplexing technology as it is the only reagent compatible with 5’ gene expression analysis. Samples were washed thoroughly with a cell staining buffer (1X PBS with 0.5% BSA), and Fc blocked for 10 min (Human TruStain FcX, Biolegend), and incubated with a cocktail containing CD3 (SP34, BD Pharmingen), 0.5 ug each of oligo tagged CD4 (TotalSeqTM-C0072, Biolegend), CD8 (TotalSeqTM-C0046, Biolegend), CCR7 (TotalSeqTM-C0148, Biolegend), CD45RA (TotalSeqTM-C0063, Biolegend), CD69 (TotalSeqTM-C0146, Biolegend), CD103 (TotalSeqTM-C0145, Biolegend), PD-1(TotalSeqTM-C0088, Biolegend), CD25 (TotalSeqTMC0085, Biolegend), and a unique hashing antibody (TotalSeqTM-C0251, C0254, C0256, or C0260, Biolegend) for an additional 30 min at 4°C. Samples were washed four times with 1X PBS (serum and azide-free), filtered using Flowmi 1000 uL pipette strainers (SP Bel-Art, Wayne, NJ), and resuspended in 300 uL FACS buffer.

CD3+ T-cells were sorted on the BD FACS Aria Fusion into RPMI (supplemented with 30% FBS). Sorted cells were counted in triplicates on a TC20 Automated Cell Counter (BioRad), washed, and resuspended in PBS with 0.04% BSA in a final concentration of 1500 cells/uL. Single-cell suspensions were then immediately loaded on the 10x Genomics Chromium Controller with a loading target of 20,000 cells. Libraries were generated using the 5’ V2 chemistry (for gene expression) and Single Cell 5□ Feature Barcode Library Kit per manufacturer’s instructions (10x Genomics, Pleasanton CA). Libraries were sequenced on Illumina NovaSeq 6000 with a sequencing target of 30,000 gene expression reads and 10,000 feature barcode reads per cell.

### 5’ Single cell RNA-Seq data analysis

For 5’ gene expression, alignments were performed using the feature and vdj option in Cell Ranger. Following alignment, hashing (HTO) and cell surface features (Antibody Capture) from feature barcoding alignments were manually updated in cellranger generated features file. Doublets were then removed in Seurat using the HTODemux function, which assigned sample identity to every cell in the matrix. Droplets with ambient RNA (cells with fewer than 400 detected genes) and dying cells (cells with more than 20% total mitochondrial gene expression) were excluded during initial QC. Data normalization and variance stabilization were performed on the integrated object using the NormalizeData and ScaleData functions in Seurat where a regularized negative binomial regression corrected for differential effects of mitochondrial and ribosomal gene expression levels. Dimensional reduction was performed using the RunPCA function to obtain the first 30 principal components and clusters visualized using Seurat’s RunUMAP function. Cell types were assigned to individual clusters using FindMarkers function with a log2 fold change cutoff of at least 0.4 (FDR ≤ 0.05) and using a known catalog of well-characterized scRNA markers for human PBMC and decidual leukocytes. 5’ feature barcoding reads were normalized using centered logratio (CLR) transformation. Differential markers between clusters were then detected using FindMarkers function. A combination of gene expression and protein markers was used to define T-cell subsets. CCR7 staining did not exhibit a significant positive peak, likely due to low signal in freshly thawed cells and hence was excluded from all downstream analyses. Only non-naïve T-cell clusters (clusters 1, 4, 5, 7, 9, 10, 13, 5, 21, 23, 24, 26, 27, Supp Figure 3C) based on relative gene expression of SELL, IL7R, and CCR7) and protein expression of CD45RA were retained for downstream analyses.

### scTCR analyses

TCR reads were aligned to VDJ-GRCh38 ensembl reference using Cell Ranger 6.0 (10x Genomics) generating sequences and annotations such as gene usage, clonotype frequency, and cell-specific barcode information. Only cells with one productive alpha and one productive beta chain were retained for downstream analyses. CDR3 sequences were required to have a length between 5 and 27 amino acids, start with a C, and not contain a stop codon. Clonal assignments from cellranger were used to perform all downstream analyses using the R package immunarch. Data were first parsed through repLoad function in immunarch, and clonality was examined using repExplore function. Diversity estimates (Chao Diversity) were calculated using repDiversity function.

### Statistical analyses

All statistical analyses were conducted in Prism 8 (GraphPad). All definitive outliers in two-way and four-way comparisons were identified using ROUT analysis (Q=0.1%) after testing for normality. Data was then tested for normality using Shapiro-Wilk test (alpha=0.05). If data were normally distributed across all groups, differences were tested using ordinary one-way ANOVA with unmatched samples. Multiple comparisons were corrected using Holm-Sidak test adjusting the family-wise significance and confidence level at 0.05. If the gaussian assumption was not satisfied, differences were tested using the Kruskall-Wallis test (alpha=0.05) followed by Dunn’s multiple hypothesis correction tests. Differences in normally distributed two groups were tested using an unpaired t-test with Welch’s correction (assuming different standard deviations). Two group comparisons that failed normality tests were tested for differences using Mann-Whitney test. For subsets of cells within a sample, differences were tested using paired t-tests.

## Results

### The immune cell landscape of maternal placental decidua at term

While the phenotypic diversity of human immune cells at the maternal-fetal interface during the first trimester has recently been described ^9^, the immunological landscape at term is less understood. Here, we characterized CD45+ decidual leukocytes and matched peripheral blood mononuclear cells (PBMC) obtained from pregnant women undergoing cesarean using a combination of single cell sequencing, flow cytometry, and functional assays (Figure 1A). Live CD45+ decidual leukocytes and PBMC collected at term from 4 subjects were sorted and profiled using single cell RNA-Seq (Figure 1A). Major decidual immune cells subsets were identified following dimension reduction, and removal of potential blood contaminants following integration of immune cells from blood and decidua (Supp Figures 1A and 1B). The expression of structural genes defining tissue-resident macrophages (VIM) and T-cells (ANXA1, EZR, TUBA1A) was higher in decidual leukocytes compared to PBMC (Supp Figure 1C), confirming their tissue resident identity.

**Figure 1:**
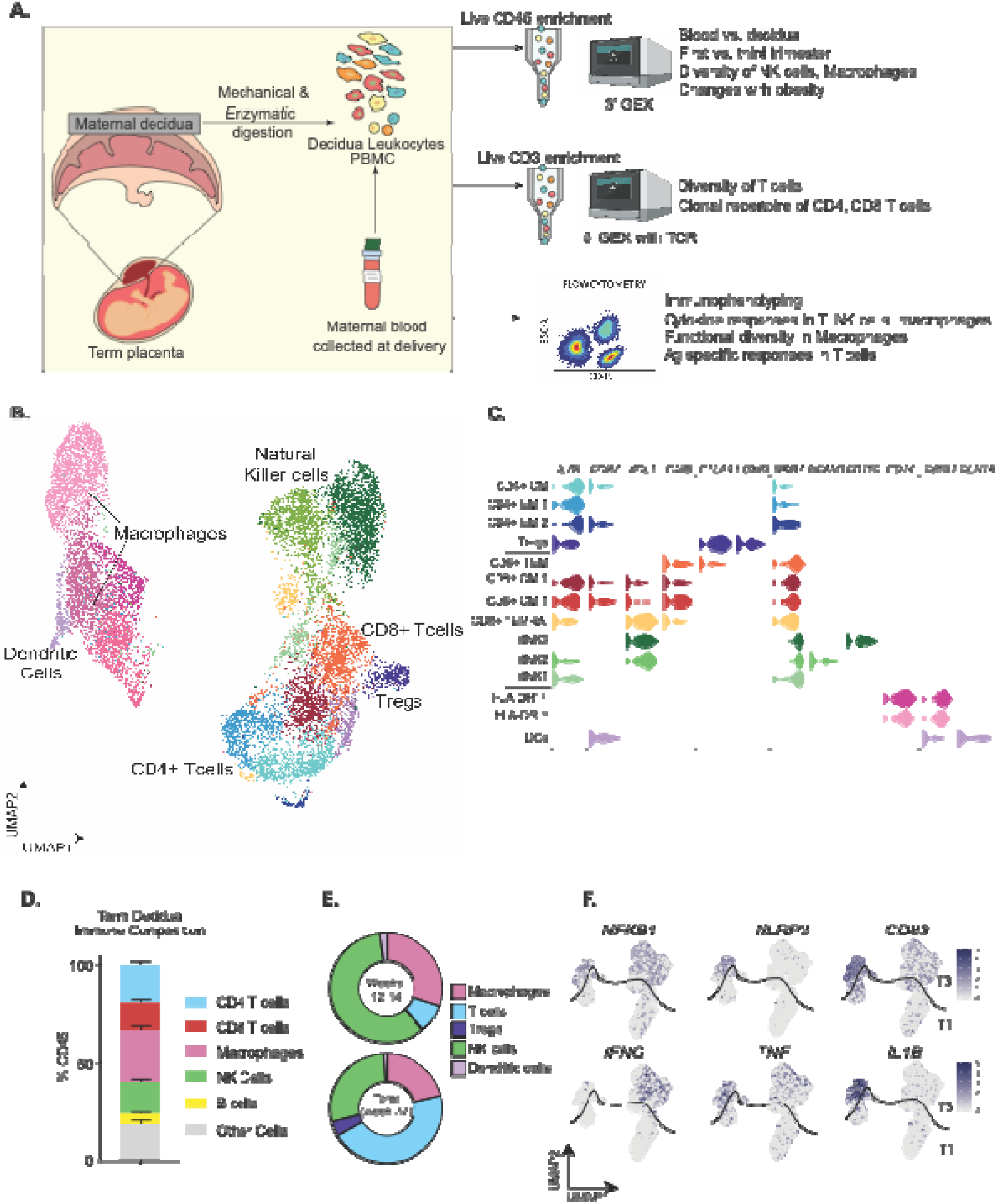
Defining the term immune landscape of placental decidua. (A) Experimental Design – CD45+ leukocytes were isolated from maternal blood and decidual membranes from term cesarean deliveries (n=4). Single-cell suspensions of decidual leukocytes and matched PBMC were subjected to gene expression profiling using 10x 3’single-cell gene expression protocol (scRNA-Seq). Additionally, sorted decidual and peripheral T-cells were surface stained with DNA oligo-tagged antibodies and hashing antibodies and subjected to 5’ gene expression and TCR on the 10x platform (n=5). Finally, the frequency, phenotype, and function of various immune cells were assayed using flow cytometry. (B) Uniform Manifold Approximation and Projection (UMAP) of 12,698 CD45+ decidual leukocytes from 4 donors. (C) Violin plots showing log-transformed normalized expression levels of major cell-type determining markers identified using Seurat. (D) Stacked bar graph comparing frequencies of major immune cell types in term decidua (n=62) using multiparameter flow cytometry. Error bars represent the mean and standard error of the mean (SEM). (E) Changes in frequencies of major immune cell subsets in placental decidua based on scRNA-Seq data of week 14 (T1) decidua ^9^ integrated with week 37 (T3) data generated in this study. (F) Feature plots comparing expression of transcription factors, surface markers, and cytokines associated with immune activation at T3 relative to T1.

Overall, the decidual immune landscape was dominated by T-cells, natural killer cells (NK), and macrophages (Figure 1B). Memory decidual CD4 and CD8 T-cells were identified based on relative expression of IL7R, CCR7 and XCL1, while regulatory T-cells (Tregs) were identified based on high expression of CTLA4 and FOXP3) (Figures 1B and 1C). Decidual natural killer cells (dNKs) were identified based on the expression of NKG7, NCAM1, and CD160; macrophages based on the expression of CD14 and CD68; and dendritic cells (DC) based on the expression of FCER1A (Figure 1C and Supp Table 1). Decidual immune cell distribution was validated using flow cytometry in a larger cohort of samples (n=62) (Supp Figure 1D, Figure 1D).

We next integrated our data with those obtained from 14-week-old decidua to understand longitudinal immunological adaptations (Supp Figures 1E, 1F). Compared to the first trimester, the relative frequencies of dNK cells and macrophages decreased while those of T-cells, notably regulatory T-cells increased at term (Figure 1E, Supp Figure 1E and 1F). Moreover, both T-cells and NK cells at term expressed high levels of IFNG and NFKB1 (Figure 1F) while decidual macrophages expressed higher levels of NFKB1 and NLRP3, activation marker CD83, and pro-inflammatory cytokines IL1B and TNF (Figure 1F), suggesting a state of heightened immune activation at term.

### Shifts in decidual natural killer (dNK) cells in term decidua

We next investigated shifts in the dNK subsets with pregnancy. Within the dNK cell population, the frequency of dNK1 (expressing B4GALNT1 and CYP26A1), which is the largest cluster found at T1 ^9^, was reduced at T3 (Supp Figure 2A). On the other hand, the frequency of dNK3 (expressing inhibitory receptors CD160 and KLRB1) increased at T3 relative to T1 (Supp Figure 2A). Finally, dNK2, a subset of cytotoxic dNKs expressing high levels of granulysin (GNLY) and granzymes (GZMB and GZMH) (Supp Figures 2A and 2B) also decreased in proportion over the course of gestation.

We then used flow cytometry to confirm and validate these shifts in dNK subsets. The dNK subsets were identified using CD103, CD49a, and ITGB2 as previously described ^9^ (Supp Figure 2C). The frequencies of dNK2 and dNK3 were significantly higher than dNK1 at T3 (Supp Figure 2D). Expression of activating (NKG2A) and inhibitory (NKG2C) receptors as well as killer immunoglobulin receptor KIR2DL4 (an activating receptor for HLA-G expressed on extravillous trophoblasts) were highest on dNK2 followed by dNK3 and low on dNK1 (Supp Figure 2E and 2F). Finally, all dNK subsets were capable of degranulation as indicated by increased levels of CD107a in response to PMA stimulation (Supp Figure 2G).

### Phenotypic and clonal diversity of decidual T-cells at term

In contrast to circulating T-cells, decidual T-cells were predominantly effector memory phenotype (Figure 2A and Supp Figure 3A). To compare the phenotypic and clonal diversity of decidual T-cells relative to blood T-cells, we performed 5’ gene expression and T-cell repertoire analyses at the single cell level coupled with cell surface protein analysis using TotalSeq-C technology (Figure 1A, n=4/group). Dimension reduction, clustering, and integration of blood and decidual T-cells revealed 32 clusters (Supp Figure 3B). Naïve T-cell clusters were identified based on high expression of IL7R, SELL, and CCR7 (Supp Figure 3C) and excluded from further analysis. The remaining 13 non-naive T-cell clusters (red font) were reclustered and identified 17 memory T-cell clusters (Figures 2B, 2C and Supp Table 2). Interestingly, there was very little overlap between the decidual and blood compartments (Supp Figure 3D), occurring within proliferating, activated CD8, and CM CD4 T-cell clusters (Figures 2B and Supp Figure 3D). Overall, decidual T-cells expressed higher levels of CD69 and PD-1 (both at protein and gene expression levels) but lower levels of SELL (encoding CD62L) compared to blood memory T-cells (Figure 2C-E and Supp Figure 3E). Single cell analyses also revealed an increased abundance of regulatory T-cells (expressing high levels of FOXP3 and CTLA4 and surface CD25) (Figure 2C, Supp Figure 3F), which was confirmed by flow cytometry (Figures 2F).

**Figure 2:**
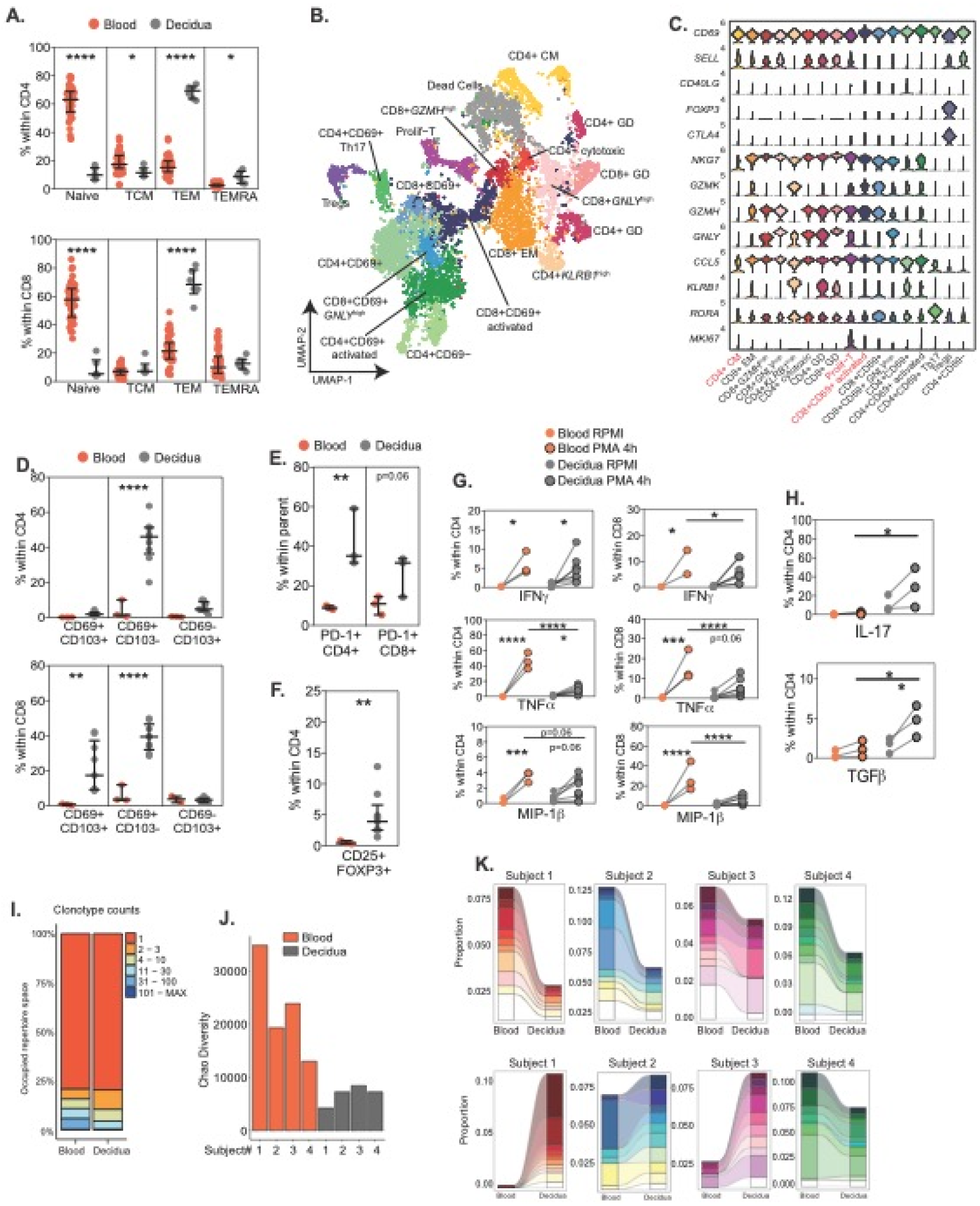
Phenotypic and clonal diversity of decidual T-cells at term. (A) Frequencies of naïve and memory CD4 and CD8 T-cell subsets in blood (n=55) and decidua (n=6) at T3 using flow cytometry. (B) UMAP of matched decidual and blood memory T-cells at term (n=4), annotated by surface- and gene-expression markers. (C) Stacked violin plots of key gene markers of memory T-cell subsets within clusters using normalized transcript counts. Red font indicated clusters shared between blood and decidual leukocytes. (D) Frequencies of CD4 (top) and CD8 T-cells (bottom) in blood (n=3) and decidua (n=9) expressing CD69 and CD103. (E-F) Frequencies of (E) PD-1+ CD4 and CD8 T-cells (n=3/group) and (F) Tregs within CD4 (n=3 for blood and n=9 for decidua). (G) Th1 cytokine responses of blood (n=3) vs decidual (n=9) CD4 (left) and CD8 (right) T-cells in response to 4h PMA/Ionomycin stimulation. (H) Treg and Th17 cytokine responses of blood (n=3) vs decidual (n=3) CD4 T-cells following 4h PMA/Ionomycin stimulation. (I) T-cell clone sizes in blood and decidua at term. (J) Chao diversity of T-cell clones from blood and decidua within each donor. (K) Clonal tracking of the 10 most abundant blood T-cell clones in the decidua (top panel) and blood (bottom panel) in each donor.

Interestingly, gene expression profiles suggest heightened activation of decidual T-cells relative to those in circulation as indicated by increased expression of NFKB1, IFNG, and TNF (Supp Figure 3G). However, Th1 cytokine production (TNFa, MIP-1b, and IFNg) by decidual T-cells following PMA/ionomycin stimulation was attenuated compared to circulating T-cells (Figure 2G). Single cell analyses also identified a Th17 (expressing RORA) cluster in the decidua (Figures 2B and 2C), which aligned with significantly higher IL-17 production by decidual compared to circulating T-cells (Figure 2H). Finally, enhanced TGF-b production following PMA/ionomycin stimulation correlated with higher frequencies of regulatory T-cells (Figure 2H).

We next compared TCR repertoire diversity between blood and decidual memory T-cells based on paired TRͰ and TRß genes captured in the single cell assay (Figure 2B). Analysis of clonal sizes indicates reduced TCR repertoire diversity in the decidua as indicated by the absence of large T-cell clones (Figure 2I) and reduced Chao diversity (Figure 2J). Additionally, in all four donors, T-cell clones that were predominant in the blood were detected in the decidua, albeit at lower proportions (Figure 2K, top panel). On the other hand, predominant decidual T-cell clones were only detected in the blood of three of the four donors (Figure 2K, bottom panel), with proportions higher in the blood in only one of the four donors.

### Two distinct subsets of macrophages exist in term maternal decidua

UMAP visualization of CD45+ cells in maternal decidua revealed two major subsets of macrophages based on surface expression of HLA-DR and CD11c (Figures 3A and 3B) as well as size/granularity (Supp Figure 4A). Both subsets had distinct transcriptional profiles compared to matched peripheral monocytes (Figure 3A). The subset expressing lower levels of HLA-DR (HLA-DR^low^) expressed higher levels of alarmins S100A8, S100A9, S100A10, IL1B, CXCL8, and M1-like associated marker TREM1 (Figures 3A and 3B). On the other hand, the subset expressing higher levels of HLA-DR (HLA-DR^high^) expressed higher levels of tetraspanins (CD9 and CD81) and canonical M2-like macrophages markers (TREM2, MSR1, APOE) (Figures 3A and 3B). The presence of these two decidual macrophage subsets was confirmed by flow cytometry (Figure 3C) and imaging flow cytometry (Figure 3D) with HLA-DR^high^ subset expressing higher levels of CD11c compared to HLA-DR^low^ subset.

**Figure 3:**
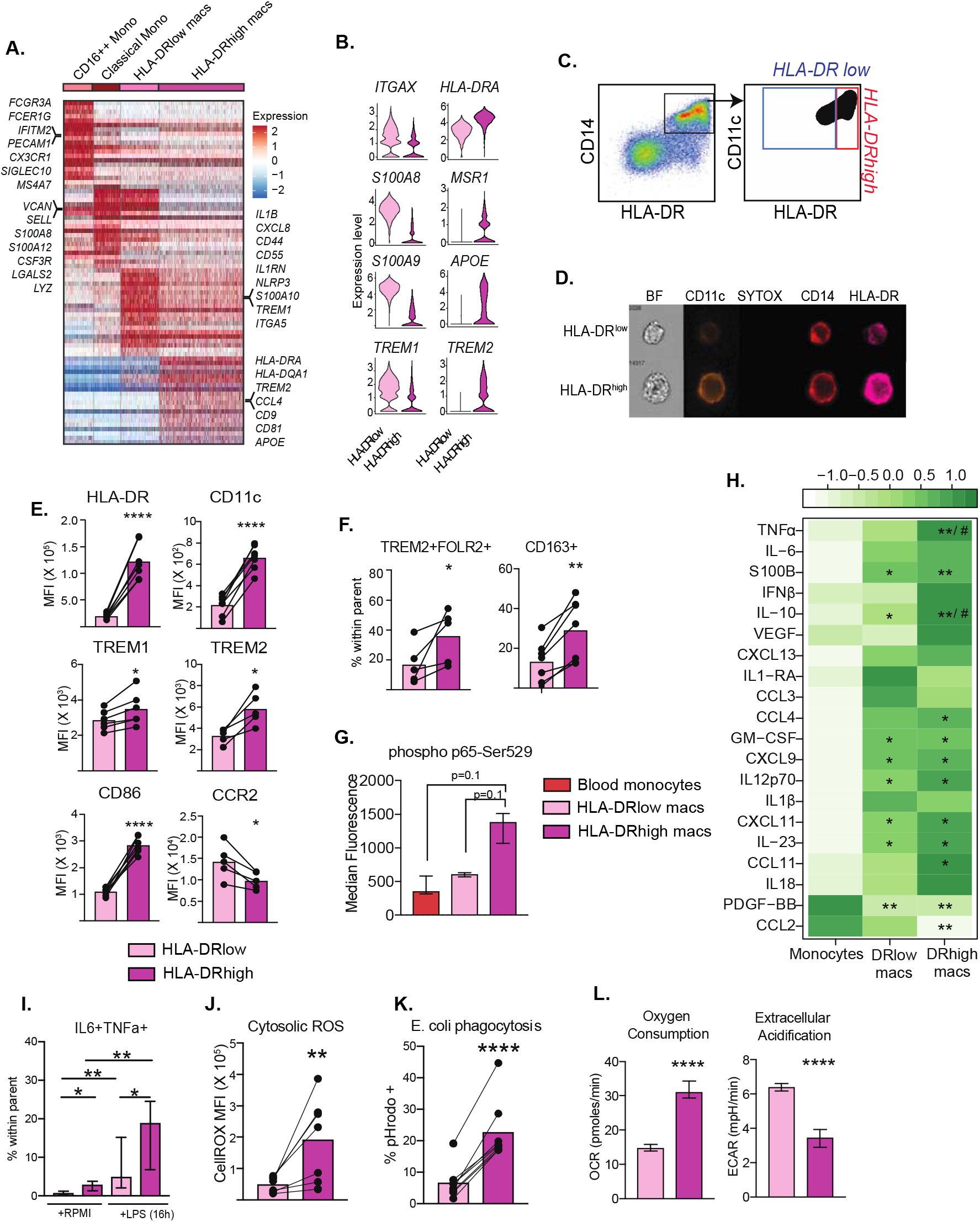
Two distinct subsets of macrophages exist in term decidua. (A)Heatmap of top marker genes for the decidual HLA-DR^low^ and HLA-DR^high^ subsets, classical and non-classical blood monocytes. Each column is a bin of a fixed number of cells and each row represents a gene. A subset of genes within each cluster is annotated. High and low expression are represented as red and blue respectively. (B) Violin plots comparing expression of gene markers within decidual macrophage subsets (C) Surface expression of HLA-DR and CD11c measured by flow cytometry. (D) Brightfield and fluorescence profiles of macrophage subsets captured using imaging flow cytometry. (E) Median Fluorescence Intensities (MFI) of macrophage surface markers on decidual macrophage subsets (n=7/group). (F) Frequencies of TREM2+ FOLR2+ (n=5/group) and CD163+ (n=7/group) within decidual macrophages. (G) Phospho p65 signal in blood monocytes and decidual macrophage subsets (n=3/group) by flow cytometry. (H) Comparison of baseline secreted cytokine profiles (median) of sorted decidual macrophage subsets (n=4/group) and blood monocytes (n=2). * and # represents significant differences relative to blood monocytes between HLA-DR^low^ and HLA-DR^high^ macrophages, respectively. (I) Responses to overnight LPS stimulation of total decidual leukocytes (n=8/group). (J-K) MFI of intracellular (J) CellROX (n=7/group) and (K) E. coli pHrodo+ (n=8/group) in decidual macrophages. (L) Oxygen consumption rate and extracellular acidification rates of decidual macrophages (n=6/group).

To further characterize these decidual macrophage subsets, we measured protein expression of several M1- and M2-like markers. Interestingly, surface expression of TREM-1 was lower on HLA-DR^low^ subset despite the higher transcript levels. Moreover, this subset expressed higher levels of surface CCR2 compared to HLA-DR^high^ subset (Figure 3E), suggesting that this population may likely be infiltrating monocyte-derived macrophages. On the other hand, HLA-DR^high^ macrophages expressed higher levels of surface markers CD86, CD11c, CD16, CD64, and TLR4 (Figure 3E, Supp Figure 4B) that are traditionally associated with M1-like macrophages. Interestingly, this subset was also enriched for regulatory CD163+ and TREM2+FOLR2+ macrophages (Figure 3E-F and Supp Figure 4C), suggesting that this population may be a tissue-resident population.

### Functional diversity of macrophage subsets in term maternal decidua

We next investigated if the transcriptional heterogeneity of decidual macrophages correlated with diversity in functional states. Flow analyses revealed that the HLA-DR^high^ macrophage subset exhibited a more heightened state of activation, expressing higher levels of phospho p65 (Figure 3G). This heightened activated state was further evident when we assessed their baseline cytokine/chemokine secretion profiles. Sorted blood monocytes, and both macrophage subsets from the same donors were cultured in RPMI for 16 hours, and their secretomes were analyzed using Luminex. Decidual macrophages secreted higher IL-12p70, IL-23, IL-10, CXCL9, S100B, GM-CSF and CXCL11 compared to blood monocytes (Figure 3H). However, secreted levels of both TNFa and IL-10 were significantly higher in HLA-DR^high^ relative to HLA-DR^low^ macrophages (Figure 3H), suggesting differential activation states between the two subsets.

Moreover, a higher percentage of HLA-DR^high^ macrophages responded to LPS stimulation as evidenced by the frequency of IL-6+TNFa+ producing cells (Figure 3I), expressed higher levels of ROS (Figure 3J and Supp Figure 4D), and were more phagocytic (Figure 3K). Since these functions are intricately linked to the metabolic state of the cell, we next compared several aspects of cellular metabolism. Our analysis shows that HLA-DR^high^ macrophages are more lipid-laden than HLA-DR^low^ macrophages (Supp Figure 4E). Moreover, seahorse energy phenotype assay indicated that at baseline, the HLA-DR^high^ macrophages had a higher oxygen consumption rate (OCR) and lower extracellular acidification rate (ECAR) (Figure 3L) correlating with higher mitochondrial membrane potential (Supp Figure 4F). Finally, HLA-DR^high^ macrophages exhibited greater uptake of glucose analog 2-NBDG and increased expression of glucose transporter GLUT1 (Supp Figure 4G). Collectively these observations suggest HLA-DR^high^ macrophages to be more metabolically active than HLA-DR^low^ cells.

### Identification of heterogeneous macrophage subsets within HLA-DR^high^

Given the elevated expression of both inflammatory (TREM1) and regulatory markers (TREM2, CD163, FOLR2) within HLA-DR^high^ macrophages (Figure 3E-F), we next investigated the heterogeneity of this subset. Higher resolution clustering and visualization of UMAP revealed 3 distinct macrophage clusters within HLA-DR^high^ cells (Figure 4A) that were transcriptionally organized along a pseudo temporal trajectory originating from HLA-DR^low^ to cluster 1, cluster 2, and terminating in cluster 3 of HLA-DR^high^ macrophages (Figure 4A and 4B). The genes defining the origin of this trajectory were primarily proinflammatory LYZ, NLRP3, TREM1, and S100 alarmins, which were highly expressed in cluster 1 (Figures 4C-D). Cells in cluster 2 expressed high levels of HLA-DR, scavenger receptors (CD36, MARCO, MSR1), lipid signaling molecules (FABP5, CD9, CD63), and regulatory molecules (TREM2 and IL1RN) (Figures 4C-D). Finally, cells in cluster 3 express high levels of complement proteins (C1QA, C1QB), folate receptor FOLR2, and negative regulators of inflammation (ATF3, KLF2, KLF4) (Figures 4C-D). We then compared the transcriptomes of these clusters to recently published data of macrophages stimulated with TPP (TNF, Pam3CSK4, and prostaglandins PGE2), a model for chronic inflammation ^38^. In concordance with the trajectory analysis and the shift from pro-inflammatory to regulatory gene expression patterns, cluster 1 of HLA-DR^high^ and HLA-DR^low^ macrophages had the highest scores while cluster 3 had the lowest score (Figure 4E).

**Figure 4:**
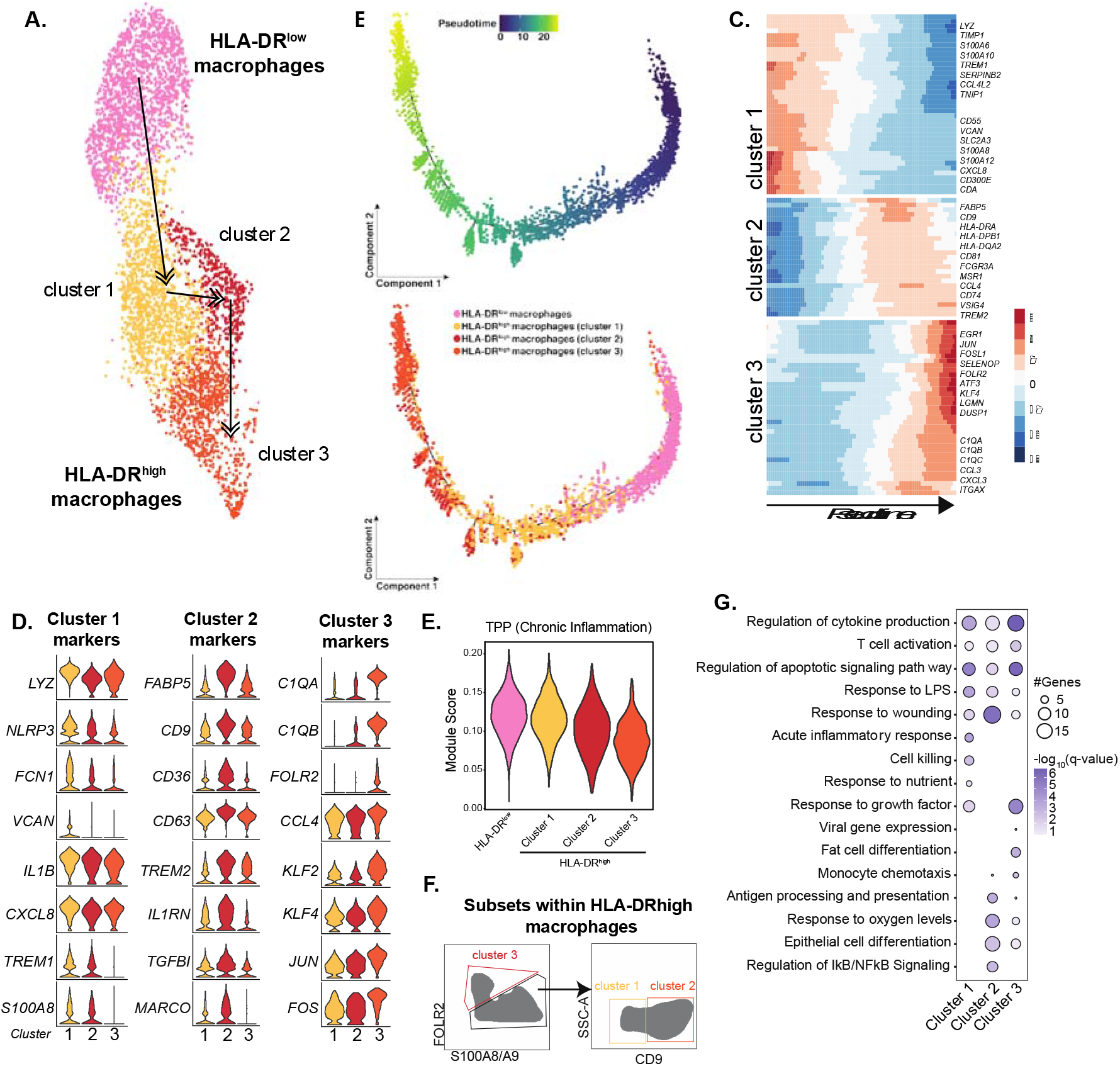
Diverse subsets within HLA-DR^high^ macrophages. (A) UMAP highlighting diverse subsets of macrophages within HLA-DR^high^ macrophages when analyzed under a higher resolution. Arrows point in the direction of pseudo temporal trajectory predicted by Monocle. (B) Trajectory of cells ordered by predicted pseudo time colored by pseudotime (top) or by the cluster of origin (bottom). (C) Heatmap of top 100 genes that define the trajectory originating from HLA-DR^low^ macrophages, through clusters 1, 2, and 3 of HLA-DR^high^ macrophages. Low and high gene expression is indicated by blue and red respectively. (D) Violin plots comparing the expression of top 8 markers from each of the HLA-DR^high^ macrophages across all three clusters. (E) Violin plot comparing aggregate score for modules of genes highly expressed in macrophages stimulated with TPP (TNF+Pam3CSK4+PGE2), an in vitro model for chronic inflammation. (F) Gating strategy for flow cytometry-based identification of HLA-DR^high^ macrophage clusters based on markers discovered from differential gene analyses. (G) Three-way functional enrichment of top genes expressed by each HLA-DR^high^ subset performed in Metascape. The size of the bubble denotes the number of genes mapping to each gene ontology (GO) term, and the intensity of color denotes the statistical strength of prediction.

Canonical markers that can differentiate the three HLA-DR^high^ sub-clusters were validated using intracellular staining of S100A8/A9, and surface expression of FOLR2 and CD9 by flow cytometry (Figure 4F). Functional enrichment of the top 100 marker genes from each sub-cluster showed that cluster1 has a dominant signature associated with LPS response, while cluster 2 had features associated with wound healing and antigen presentation, and finally cluster 3 was enriched in regulatory pathways (Figure 4G).

### Maternal obesity is associated with reduced frequency of decidual T-cells and functional rewiring of HLA-DR^high^ decidual macrophages

Pre-pregnancy (pregravid) obesity is increasing around the world and is implicated in elevated risk for perinatal complications which may have long-term impact on maternal and offspring health. Therefore, we next investigated the impact of maternal pregravid obesity on the immunological landscape of the decidua. Frequency of major decidual immune subsets from lean pregnant women (BMI <25) was compared to those with obesity (BMI>30). Obesity had no impact on dNK cell frequencies (Figure 5A). In contrast, the proportions of CD4 and CD8 T-cells within the CD45 compartment were significantly reduced (Figure 5A). Moreover, frequencies of both decidual CD4 and CD8 T-cells negatively correlated with maternal pregravid BMI (Supp Figures 5A). However, no differences in the relative abundance of naïve/memory T-cell subsets (Supp Figure 5B) or tissue-resident subsets (Supp Figure 5C) were observed with pregravid obesity. Furthermore, T-cell responses to PMA/ionomycin stimulation were comparable between the two groups (Supp Figure 5D).

**Figure 5:**
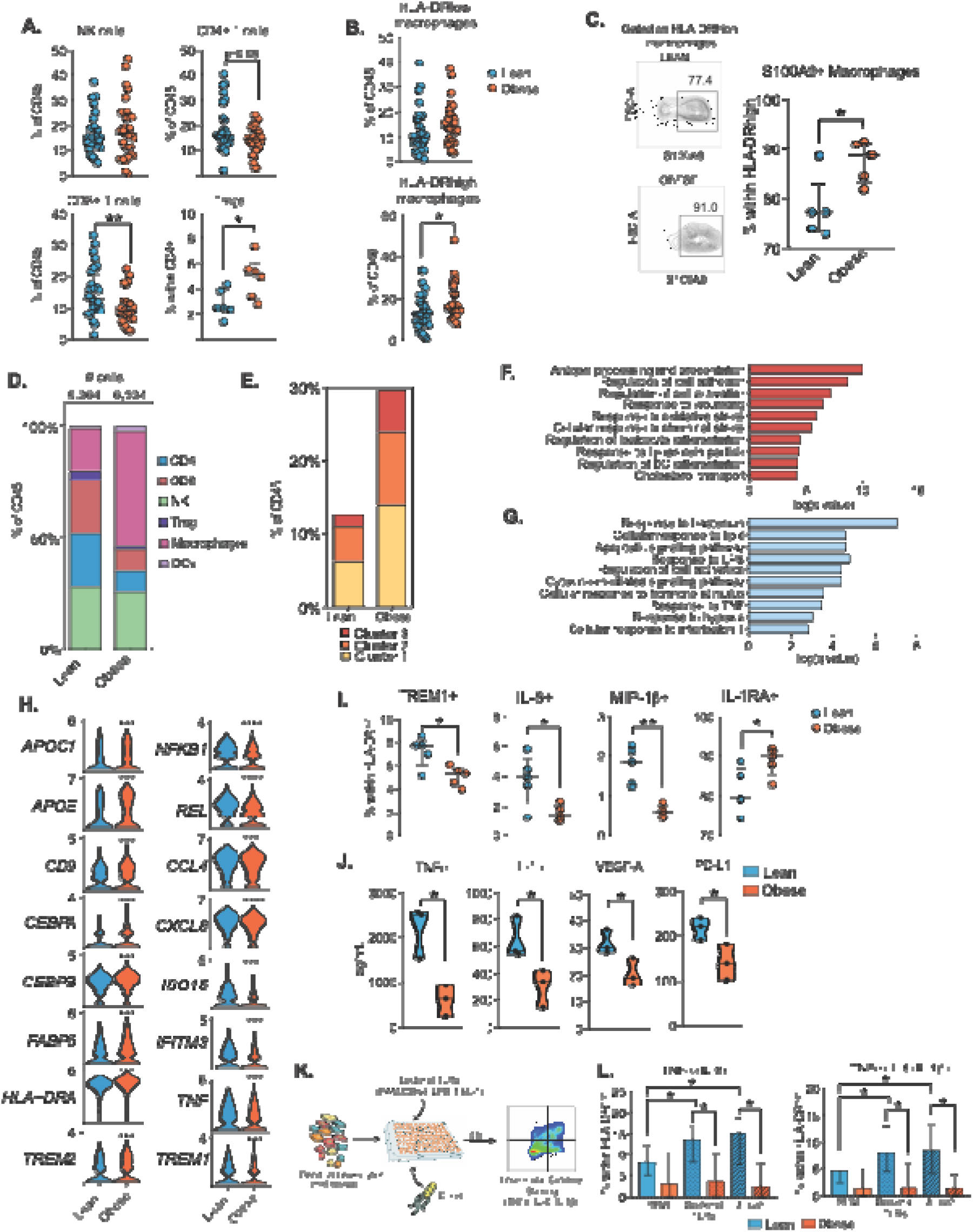
Decidual immune cell composition with maternal obesity. (A-B) Dot plots comparing (A) lymphocyte and (B) macrophage proportions within CD45+ decidual cells in lean pregnant women (n=6 for Tregs, 32 for rest) and those with obesity (n=6 for Tregs, 31 for rest). (C) Gating strategy and dot plots comparing frequencies of S100A9+ cells within HLA-DR^high^ macrophages (n=5/group). (D) Stacked bar graph comparing frequencies of decidual leukocytes within total cells sequenced in each group (within CD45+ cells). (E) Stacked bar comparing HLA-DR^high^ macrophage cluster within total CD45+ cells. (F-G) Gene Ontology (GO) terms of genes that are up- (F) and down- (G) regulated with pregravid obesity within HLA-DR^high^ macrophages predicted by Metascape. Only differentially expressed genes with a Log_2_ fold change cutoff of 0.4 were analyzed. (H) Violin plots compared normalized transcripts of select genes that are significantly up- (left) or down- (right) regulated with obesity. (I) Dot plots comparing frequencies of TREM1+ cells or cytokine expressing cells within HLA-DR^high^ macrophages (n=5/group). (J) Violin plots comparing secreted factors at baseline by FACS sorted HLA-DR^high^ macrophages (n=3/group). (K) Experimental design for measuring ex vivo macrophage responses to E. coli or bacterial TLRs. (L) Bar graphs comparing cytokine-producing cells TNFa+IL6+ (left) and TNFa+IL6+IL1b+ (right) following stimulation with bacterial TLRs and E. coli (n=6/group).

The reduction in the frequency of decidual T-cells with pregravid obesity was accompanied by an expansion of decidual macrophages (Supp Figure 5E), driven by the HLA-DR^high^ subset (Figure 5B). Moreover, the frequency of HLA-DR^high^ but not HLA-DR^low^ subsets positively correlated with maternal pregravid BMI (Supp Figure 5F). Flow analyses of HLA-DR^high^ macrophages revealed an increase in S100A9+ macrophages (Figure 5C), a marker of subclusters 1 and 2 (Figure 4F), and recently infiltrated macrophages ^39^.

Next, we leveraged scRNA-Seq to further interrogate the impact of pregravid obesity on the transcriptional landscape of decidual leukocytes (Supp Figure 6A). In line with the flow cytometry data, this analysis revealed a contraction of CD4 and CD8 T-cells and expansion of decidual macrophages (Figure 5D), driven by increased frequencies of all three HLA-DR^high^ subsets (Figure 5E). Differential gene expression analyses of HLA-DR^high^ macrophages between lean and obese groups revealed an up-regulation of pathways associated with response to lipoproteins and cholesterol transport (FABP5, CD36, APOE, APOC1, CD9), response to stress (CEBPA, CEBPB, HSPA5, HSPA8), and antigen processing and presentation (HLA-DRA) (Figures 5F, 5H left column, Supp Figure 6B left column, and Supp Table 3). Proinflammatory signatures, on the other hand, were downregulated with pregravid obesity (Figure 5G and Supp Table 3). This included cytokine signaling pathways (CCL4, CXCL8, TNF, NFKB1, REL), interferon-stimulated genes (ISG15, IFITM3), metabolic genes (COX6A1, COX6B1), and genes important for T-cell activation (CD55, CD300E) (Figures 5G, 5H right column, and Supp Figure 6B right column).

To validate the transcriptional changes described above, we assessed the protein expression of inflammatory mediators and markers using flow cytometry. While frequencies of TREM1+, MIP-1b+, and CXCL8+ decidual macrophages were reduced with pregravid obesity (Figure 5I), the frequency of IL-1RA+ macrophages increased (Figure 5I). Additionally, spontaneous production of several proinflammatory cytokines (TNFa, IL-1b, IL-12p70) was significantly attenuated within sorted HLA-DR^high^ macrophages (Figure 5J and Supp Figure 6C) in the absence of changes in baseline levels of activation-associated signaling molecules (Supp Figure 6D). To define the biological significance of the downregulation of genes important for anti-microbial function with pregravid obesity (Figure 5G), we measured the frequency of cytokine producing decidual macrophages following stimulation with a cocktail of bacterial TLRs ligands (activating TLRs 1/2/4/6) or E. coli using flow cytometry (Figure 5K and Supp Figure 6E). Pregravid obesity was associated with reduced frequency of TNF+IL-6+IL-1b+ macrophages following both stimulation conditions within both HLA-DR^high^ (Figure 5L) and HLA-DR^low^ subsets (Supp Figure 6E).

## DISCUSSION

Roughly 30-40% of maternal membranes of human placenta (or decidua) are immune cells that play a critical role in the maintenance of fetomaternal homeostasis ^40 2^. The phenotype and frequencies of immune cells in the maternal decidua dynamically adapt to fetal needs, mediating fetal growth and tolerance, protection from infections, and ultimately facilitating parturition ^40, 41^. Recent advances in single cell technologies have facilitated the unbiased profiling of rare populations of immune cells residing in the early placental decidua ^9, 42^. While the transcriptional profiles of placental trophoblast populations from fetal membranes under homeostasis ^43, 44^ and in preterm labor have been recently investigated ^45^, an atlas of immune diversity of human decidua at term is still lacking. The availability of such a high-resolution map of immune cells collected from individuals with no pregnancy complications would pave the way for future research on the role of decidual T, NK cell, and macrophage in the pathophysiology of adverse outcomes such as preeclampsia, spontaneous abortion, and preterm labor. We, therefore, profiled the immune compartment of term decidual membranes, isolated from mothers undergoing scheduled cesareans to generate a detailed map of transcriptomic, phenotypic, and functional diversity of term decidual T-cells, NK cells, and macrophages. Finally, given the strong association between maternal pre-pregnancy BMI and pregnancy complications ^3–7^, we investigated the impact of pregravid obesity on the functional rewiring of decidual immune cells.

The early placental decidua is populated by NK cells (dNKs) ^46, 47^, which account for about 70% of decidual leukocytes that are recruited from blood by factors such as IL-15 and progesterone ^48, 49^. Early dNKs facilitate embryonic development and promote decidual tolerance to the embryo. Furthermore, animal studies have revealed the critical role of dNK-derived factors (VEGF-C, TGF-b, IFNg) in uterine vascular modifications and EVT differentiation ^50, 51^. Single cell RNA-seq and mass cytometry analyses of first-trimester decidua ^9, 10^ revealed three subpopulations of dNKs. Of the three populations, dNK1s (Eomes+) are the most abundant, expressing granzymes, KIR inhibitory receptors and LILRB1 suggesting their preferential interactions with EVTs. This subset of dNKs produce growth factors such as pleiotrophin and osteoglycin, promoting fetal growth in both mice and humans ^52^, suggesting their critical role in early pregnancy. On the other hand, mass cytometry analyses suggest that dNK3s are predominantly Tbet+ and mounted the strongest polyfunctional (CD107a, MIP-1a, MIP-1b, XCL1, IFNg) response to ex vivo PMA stimulation ^10^. Our analyses reveal a sharp drop in total dNK proportions at term, with a substantial shift towards dNK2 and dNK3 cells. This reduction is in line with the reduced role of immune cells in vascular remodeling and promotion of embryonic growth but enhanced immune activation associated with labor in the third trimester ^53^. While it is known that dNKs become less granular over the course of a healthy pregnancy ^54^, the impact of changes in dNK frequencies and activation status to labor, parturition, and adverse pregnancy outcomes associated with late-stage pregnancies is still unclear.

Reduction and redistribution of dNK cell subsets with gestation was associated with an expansion of T-cells, potentially indicating a shift from tissue remodeling to anti-microbial responses. T-cells are recruited into the decidua from the blood by trophoblast-derived CXCL16. Tissue-resident memory (TRM, CD69+) CD8 T-cells are present in term decidua, and a subset of these cells express CD103. TRM T-cells have been reported to express high levels of PD-1 but low levels of effector molecules such as GZMB and perforin compared to their blood counterparts in line with their ability to trigger trophoblast antigen-specific tolerance ^55, 56^. We also observed increased PD-1 expression on decidual CD4 T-cells, however, these TRMs were predominantly CD103-. Aberrant Th1 responses in placental CD4 T-cells have been associated with pathologies such as recurrent spontaneous abortions (RSA) ^57^. Our data indicate reduced ability of decidual CD8 T-cells to produce IFNg, MIP-1b, and TNFa compared to circulating CD8 T-cells. While decidual and circulating CD4 IFNg responses were comparable, production of MIP-1b, and TNFa were lower in decidual CD4 T-cells compared to their blood counterparts. Th17 responses were significantly higher in the decidua. Both Th1 and Th17 responses at the maternal-fetal interface have been shown to increase over the course of pregnancy and play a role in the initiation of labor ^58^. Finally, CD4 T-cells generated higher TGF-b responses in line with increased frequency of Tregs, which play an important role in maintaining and preventing miscarriages ^59^.

To our knowledge, this is the first study comparing the clonality of placental and blood T-cells from matched donors. In all four healthy donors, we observed that decidual T-cells repertoire was more restricted. However, in all four donors, the dominant decidual T-cell clones were also detected in the blood, suggesting their recruitment from the blood. The process by which these specific clones are recruited and retained in the decidua as well as their antigen specificity remains poorly understood. Studies have shown that decidual CD4 and CD8 T-cells can recognize viral antigens during Zika and dengue infections ^60^ and that frequency of EBV and CMV-specific CD8 T-cells is greater in the decidua compared to blood ^61^. These findings suggest that decidual T-cells can mediate anti-microbial responses.

Compared to decidual lymphocytes, macrophage frequencies modestly decreased at term relative to first term decidua. A large macrophage population with the near absence of DCs suggests their potential role in antigen presentation at the maternal-fetal interface. Additionally, these cells play a critical role in regulating T-cell responses, tissue remodeling, and clearance of apoptotic cells. Two unique subsets of macrophages have been defined in decidua during early gestation – characterized by differential surface expression of CD11c and CCR2 ^62^. Gene expression profiling of these subsets suggested that the CD11c^low^ subset plays a role in tissue homeostasis while CD11c^high^ are important for antigen presentation ^63^. However, neither subset conformed with the M1/M2 macrophage paradigm, suggesting an underappreciation of their plasticity at the maternal-fetal interface. Using unbiased single-cell RNA-sequencing of matched term decidual macrophages and blood monocytes, we report two major macrophage subsets characterized by co-expression of HLA-DR and CD11c as described for first trimester decidua ^63^.

Trajectory analysis, flow cytometry, and profiling of secreted cytokines at baseline revealed that the HLA-DR^low^ macrophages are an inflammatory subset that are transcriptionally similar to blood monocytes. On the other hand, despite high expression level of regulatory molecules (TREM2, CD163, CD206), HLA-DR^high^ macrophages exhibited an enhanced capacity for TLR4 responses and phagocytosis. Furthermore, HLA-DR^high^ macrophages also accumulated higher levels of lipids and are metabolically primed for oxidative phosphorylation, supporting their identity as bonafide tissue-resident macrophages, as previously described in other compartments ^64^. Further characterization of the heterogeneity within this subset revealed three sub-populations predicted to play a role in acute inflammatory responses and killing (S100A8+ CD9-subset), response to wounding (S100A8+ CD9+), and apoptosis and faT-cell differentiation (S100A8-FOLR2+). Therefore, decidual macrophages span the spectrum from inflammatory blood-derived HLA-DR^low^ cells to HLA-DR^high^ FOLR2+ regulatory cells involved in tissue remodeling and apoptosis of cellular debris.

A consistent theme across both decidual lymphocyte and macrophage populations was a heightened state of immune activation at term compared to the first trimester. This is evident from increased levels of inflammatory cytokine transcripts (IL1B, IFNG, TNF) across dNK, T-cell, and macrophage subsets and higher levels of spontaneous secretion of cytokines and chemokines by sorted macrophage subsets compared to blood monocytes. We posit that this state of activation is important for parturition ^65^ and is in line with recent studies highlighting progressive peripheral immune activation associated with gestation ^66, 67^. Understanding these pregnancy-associated shifts in immune states is therefore critical in addressing the immunological aberrancy that contributes to pregnancy complications. We have previously demonstrated that maternal obesity alters this state of activation in peripheral monocytes, resulting in dampened ex vivo cytokine responses to TLR4 stimulation ^22^. This is a critical observation given that pregravid obesity is strongly associated with placental abruptions, preeclampsia, delayed wound healing, and delivery complications including prolonged first stages labor and higher rates of cesarean sections ^22, 68–70^.

Our analyses suggest a dramatic drop in CD4 and CD8 T-cell numbers and concomitant increases in macrophage frequencies with pregravid obesity driven in a BMI-dependent manner by increased frequency of the HLA-DR^high^ macrophage subset. Despite the overall loss of T-cells, we report no preferential loss of any T-cell subset or functional loss in cytokine responses. In contrast, macrophage responses were significantly attenuated – with the downregulation of genes involved in pathogen sensing and cytokine signaling. Functionally, this translated to reduced levels of secreted inflammatory cytokines at baseline (TNFa, IL-1b) but also poor responses to ex vivo E. coli stimulation. Taken together, these findings highlight an establishment of a tolerance-like program in term decidual macrophages with maternal obesity, mirroring the phenotype observed in peripheral monocytes reported in our previous study ^22^. We posit that the skewing of term decidual microenvironment and macrophages to a regulatory phenotype is one mechanism of protecting the developing fetus from the undesirable effects of maternal inflammation. Suboptimal activation and attenuated responses can have untoward consequences for both host defense and initiation of labor. These changes could also contribute to obesity-associated pathologies during the late stages of pregnancy.

Our study is limited by restricting analyses of decidua collected following cesarean sections. Future studies need to focus on identifying key immunological changes occurring in the placenta stratified by the mode of delivery. The dietary contributions to maternal obesity-associated placental adaptations are also unclear. Future research will need to focus on establishing immunological changes during early and mid-gestation decidua to identify early dysfunction in vascularization, tissue remodeling, and fetal development in animal models of diet-induced obesity. Finally, while pregravid obesity has been shown to attenuate cord blood monocyte and T-cell responses, its influence on fetal macrophage populations in chorionic villi is still unknown. Evidence of such influence can have far-reaching consequences on placental angiogenesis, remodeling, debris clearance, and host defense.

## Author Contributions

Conceptualization, S.S., N.E.M., and I.M.; methodology, S.S., N.E.M., and I.M.; investigation, S.S., B.D., and N.M; writing, S.S. and I.M.; funding acquisition, N.E.M., and I.M.; resources, M.R., N.E.M. All authors have read and approved the final draft of the manuscript.

## Acknowledgments

We are grateful to all participants in this study. We thank the MFM Research Unit at OHSU for sample collection and Allen Jankeel, Michael Z. Zulu, Gouri Ajith, Isaac Cinco, and Hannah Debray at UCI for assistance with tissue processing. We thank Dr. Jennifer Atwood at the UCI Institute for Immunology Flow Cytometry Core for assistance with FACS sorting, imaging flow cytometry and Dr. Melanie Oakes at the UCI Genomics Research and Technology Hub (GRT Hub) for assistance with 10x library preparation and sequencing.

## Data availability

The datasets supporting the conclusions of this article are available on NCBI’s Sequence Read Archive PRJNA817521 and PRJNA887020. https://dataview.ncbi.nlm.nih.gov/object/PRJNA887020?reviewer=ffmuhfajbmccgot3bp29c7fa

