## Supplemental Figures for "Multimodal profiling of term human decidua reveals tissue-specific immune adaptations with maternal obesity"

**SUPPLEMENTARY FIGURES**

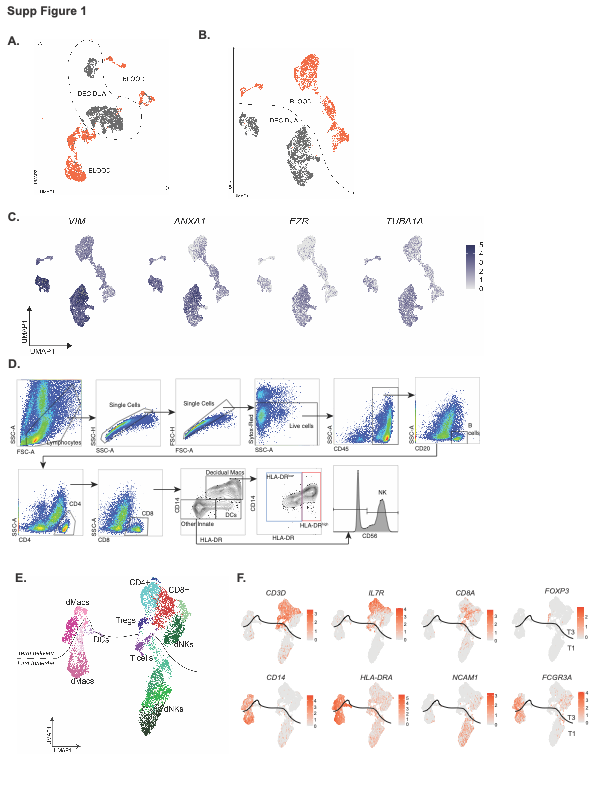

**Supplementary Figure 1: Comparison of peripheral and decidual leukocytes in early and late gestation.**

(A, B) UMAP representation of integrated blood and decidua profiles before (A) and after (B) the removal of infiltrating blood cells from decidua leukocytes. (C) Feature plots showing increased expression of tissue residency markers in decidual leukocytes compared to peripheral blood cells. (D) Gating strategy for quantification of immune cell subsets in term decidua. (E) UMAP representation of integrated decidua profiles from first (T1) ^9^ and third trimesters (T3) (downsampled to 6,116 cells each). (F) Feature plots comparing expression of key immune cell markers at T1 and T3.

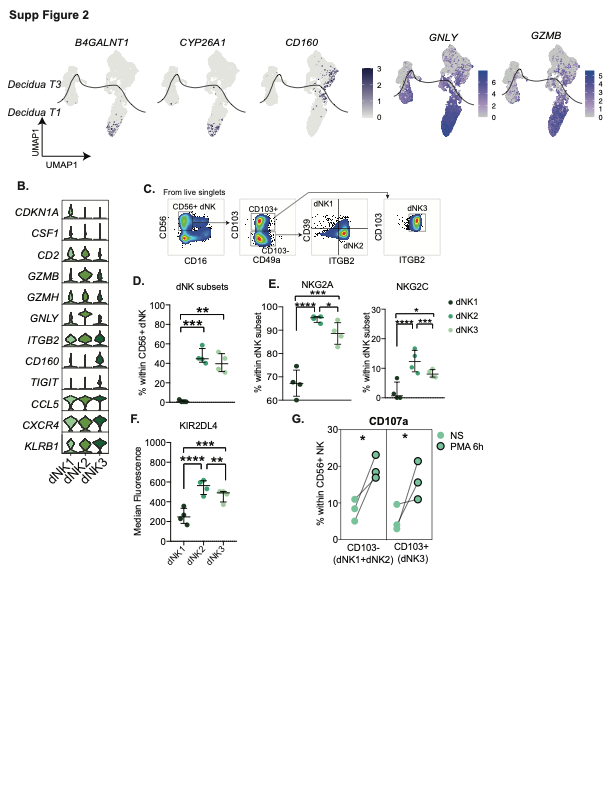

**Supplementary Figure 2: Characterization of decidual NK cell populations in term decidua**

(A) Feature plots of key dNK (decidual NK) cell markers and cytokines at T1 and T3. (B) Violin plots comparing key NK cell markers across the three decidual NK cell subsets at T3. (C) Gating strategy for identification of decidual NK subsets as previously described in the first trimester decidua ^9^. (D) Dot plots comparing proportions of dNK subsets (n=4) characterized within CD56+ NK cells in term decidua using flow cytometry. (E) Dot plots comparing frequencies of dNK subsets expressing activating and inhibitory receptors (n=4). (F) KIR2DL4 expression (HLA-G activator) across dNK subsets (n=4). (G) CD107 expression within CD103+ (dNK3) and CD103- (dNK1 and dNK2) dNK subsets (n=3) following PMA stimulation

**Supplementary Figure 3: Comparing decidual T-cells with blood T-cells**

(A) Gating strategy for identifying memory T-cells (left) and tissue-resident T-cell subsets (right) in the blood and decidua. (B) Integrated UMAP of blood and decidual CD3+ T-cells colored by cluster (left) and source (right). Non-naïve clusters are highlighted in red. (C) Violin plots comparing key T-cell markers across all clusters of CD3+ cells. Non-naïve clusters are in red font. (D) UMAP projection of memory T-cells in blood vs. decidua (n=4/group). (E) Feature plot of surface PD-1 and CD69 expression across blood and decidual memory T-cells. (F) Bubble plot comparing surface expression of protein markers across blood and decidual memory T-cell clusters. * Indicate clusters shared between blood and decidual clusters. (G) Violin plots comparing markers of activation between blood and decidual memory T-cells.

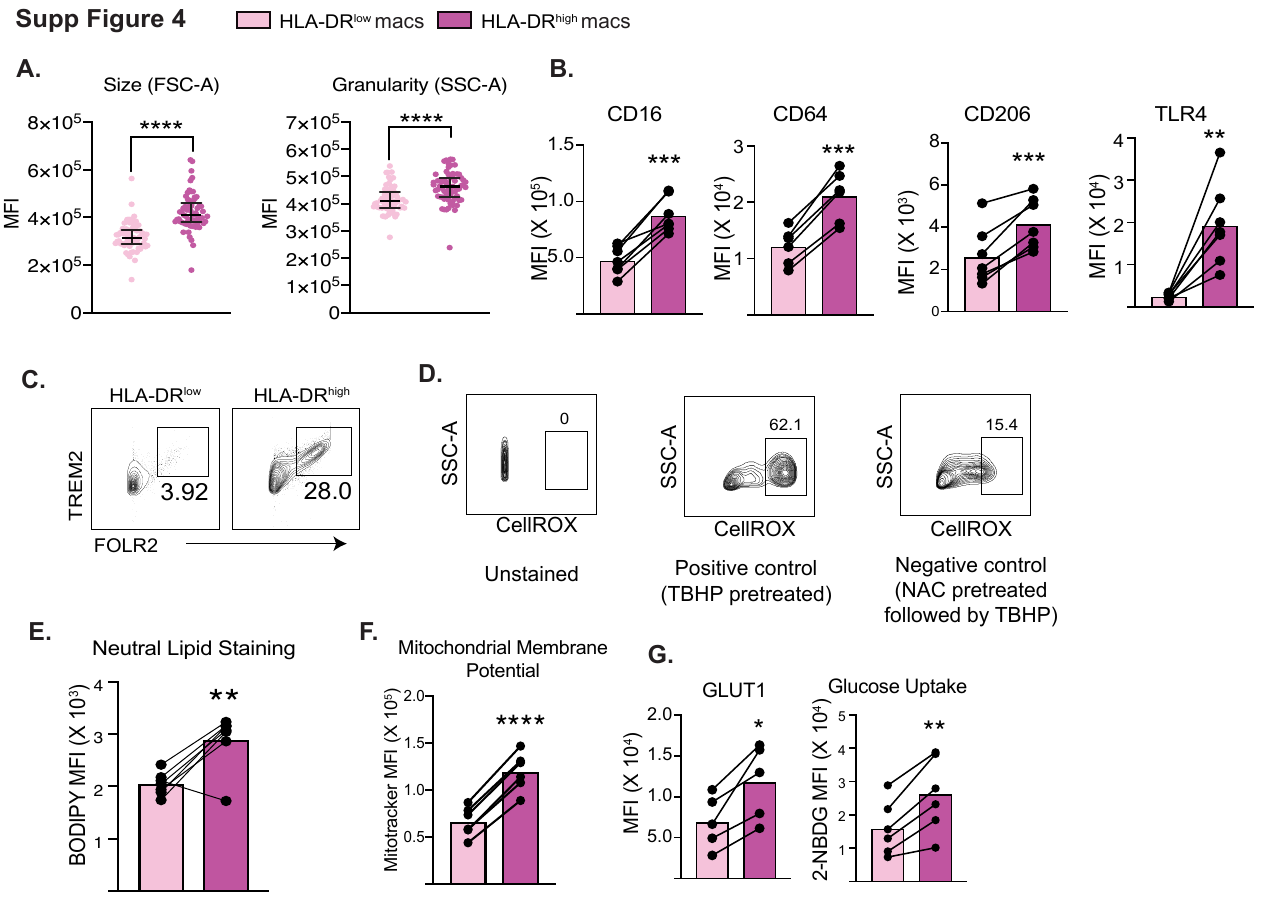

**Supplementary Figure 4: Functional differences between HLA-DR^low^ and HLA-DR^high^ macrophages**

(A) Dot plots comparing size (FSC) and granularity (SSC) of decidual macrophage subsets (n=63/group). (B) Bar graphs comparing median fluorescence of surface markers between HLA-DR^low^ and HLA-DR^high^ macrophages (n=6) (C) Representative FACS plots for identification of TREM2+ FOLR2+ macrophage populations within HLA-DR^low^ and HLA-DR^high^ macrophages (D) Representative CellROX staining within macrophages (n=6). Oxidant tert-Butyl hydroperoxide (TBHP) pretreatment served as a positive control and antioxidant pretreatment served as a negative control. (E-G) Dot plots comparing MFI of (E) BODIPY stain (n=7), (F) mitochondrial membrane potential (n=6), and (G) surface GLUT1 (n=5) and internalized 2-NBDG (n=6) between HLA-DR^low^ and HLA-DR^high^ macrophages.

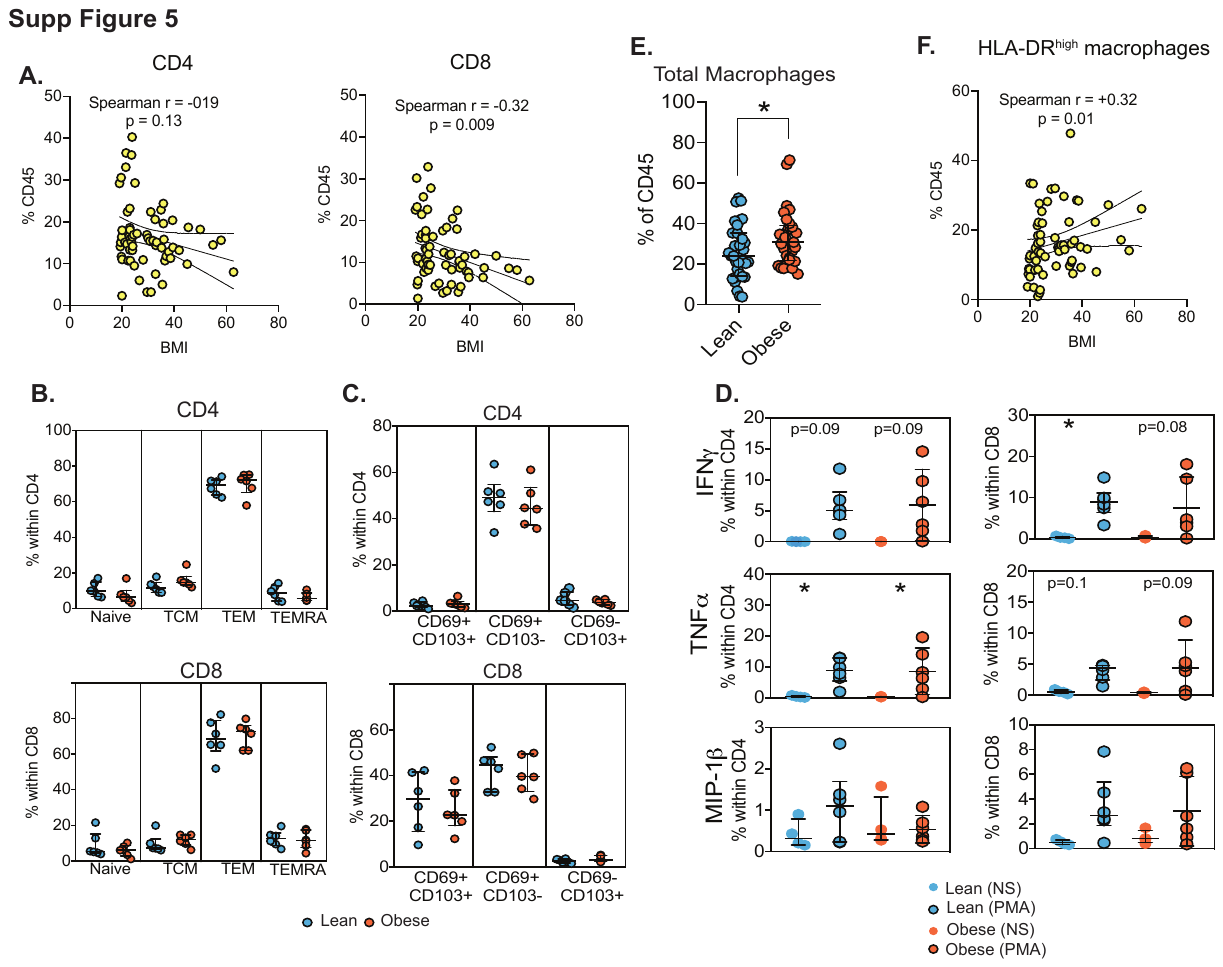

**Supplementary Figure 5: Impact of maternal obesity on decidual lymphocytes.**

(A) Correlation graphs comparing % decidual CD4 and CD8 T-cells as a function of maternal pre-pregnancy BMI (n=63). (B, C) Relative proportions of (B) naïve/memory T-cell subsets and (C) tissue-resident T-cell subsets within decidual CD4 (top) and CD8 T-cells (bottom) (n=6/group). (D) Dot plots comparing cytokine responses to PMA/ionomycin stimulation in decidual CD4 (left) and CD8 T-cells (right) (n=6/group). (E) Dot plots comparing proportions of macrophages within CD45+ cells (n=32 leans, 31 obese). (F) Correlation graphs comparing HLA-DR^low^ and HLA-DR^high^ macrophages in relation to maternal pre-pregnancy BMI (n=63).

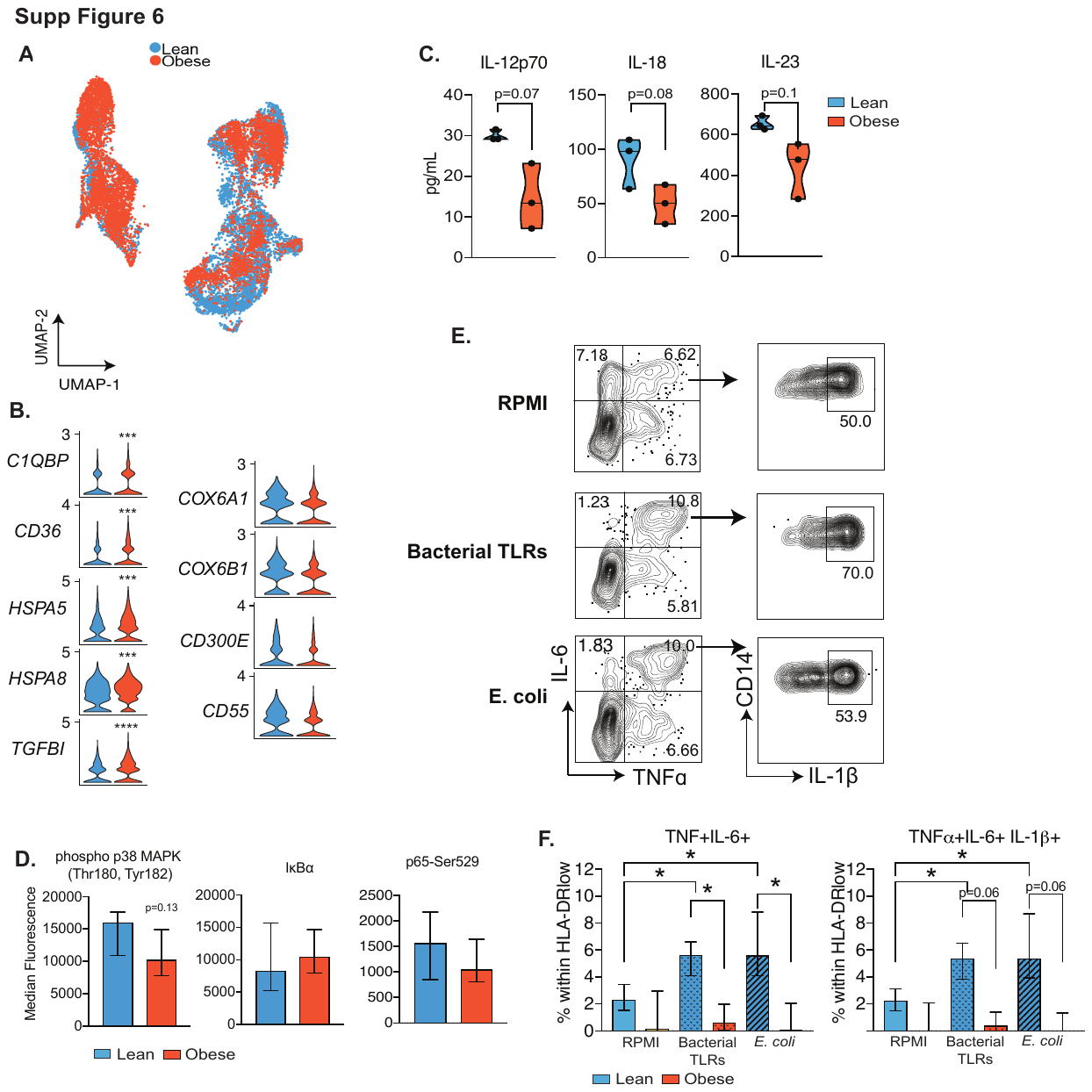

**Supplementary Figure 6: Impact of maternal obesity on decidual macrophages.**

(A) UMAP projection of CD45+ compartment within term decidual cells highlighted by contributions from lean (blue) and obese (orange) subjects (n=2/group). (B) Violin plots comparing relative expression of genes up (left) or downregulated (right) with maternal obesity. (C) Violin plots comparing secreted factors at baseline by FACS sorted HLA-DR^high^ macrophages (n=3/group). (D) Bar graphs comparing intracellular NF-kB signaling mediators downstream of TLR4 measured using flow cytometry (n=6/group). (E) Gating strategy for quantification of cytokine-producing macrophages after bacterial TLRs and E. coli stimulation. (F) Bar graphs comparing cytokine-producing cells within HLA-DR^low^ macrophages - TNFa+IL6+ (top) and TNFa+IL6+IL1b+ (bottom) following stimulation with bacterial TLRs and E. coli (n=6/group).

Supp Table 1: Gene markers for clusters identified by 3′ gene expression analysis of total decidua, related to Figure 1.

Supp Table 2: Gene markers for clusters identified by 5′ TCR/gene-expression analysis in decidua, related to Figures 2 and Supp Figure 3.

Supp Table 3: Gene markers for macrophage clusters identified by 3′ gene expression analysis, related to Figure 5.
